# Expression of novel fusion antiviral proteins Ricin A Chain-Pokeweed Antiviral Proteins (RTA-PAPs) in Escherichia coli and their inhibition of protein synthesis and of hepatitis B virus in vitro

**DOI:** 10.1101/322040

**Authors:** Yasser Hassan, Sherry Ogg, Hui Ge

## Abstract

Ricin A chain (RTA) and Pokeweed antiviral proteins (PAPs) are plant-derived N-glycosidase ribosomal-inactivating proteins (RIPs) isolated from R*icinus communis* and *Phytolacca Americana* respectively. This study was to investigate the potential antiviral value of novel fusion proteins between RTA and PAPs (RTA-PAPs). In brief, RTA-Pokeweed antiviral protein isoform 1 from seeds (RTA-PAPS1) was produced in *E. coli in vivo* expression system, purified from inclusion bodies using gel filtration chromatography and protein synthesis inhibitory activity assayed by comparison to the production of a control protein Luciferase. The antiviral activity of the RTA-PAPS1 against Hepatitis B virus (HBV) in HepAD38 cells was then determined using a dose response assay by quantifying supernatant HBV DNA compared to control virus infected HepAD38 cells. The cytotoxicity in HepAD38 cells was determined by measuring cell viability using a tetrazolium dye uptake assay. Results showed that RTA-PAPS1 could effectively be recovered and purified from inclusion bodies. The refolded protein was bioactive with 50% protein synthesis inhibitory concentration (IC50) of 0.06nM (3.63ng/ml). The results also showed that RTA-PAPS1 had a synergetic activity against HBV with a half-maximal response concentration value (EC50) of 0.03nM (1.82ng/ml) and a therapeutic index of >21818. The fusion protein was further optimized using *in silico* tools, produced in *E. coli in vivo* expression system, purified by three-step process from soluble lysate and protein synthesis inhibition activity assayed. Results showed that the optimized protein RTA mutant-Pokeweed antiviral protein isoform 1 from leaves (RTAM-PAP1) could be recovered and purified from soluble lysates with gain of function activity on protein synthesis inhibition with an IC50 of 0.03nM (1.82ng/ml). Collectively, our results demonstrate that RTA-PAPs are amenable to effective production and purification in native form, possess significant antiviral activity against HBV *in vitro* with a high therapeutic index and, thus, meriting further development as potential antiviral agents against chronic HBV infection.

## 1 Introduction

^i^Pokeweed antiviral proteins (PAPs) are expressed in several organs of the plant pokeweed (*Phytolacca Americana*) and are potent type I Ribosome Inactivating Proteins (RIPs). Their sizes vary from 29-kDa to 30-kDa and are able to inhibit translation by catalytically removing specific adenine residues from the large rRNA of the 60S subunit of eukaryotic ribosomes (Domashevskiy and Goss, 2015; Kurinov and Uckun, 2003; Rajamohan et al., 2001). Furthermore, PAPs can depurinate specific guanine residues, in addition to adenine, from the rRNA of prokaryotic ribosomes. PAPs possess antiviral activity on a wide range of plant and human viruses through various mechanisms (Domashevskiy and Goss, 2015). Transgenic plants expressing different forms of PAPs were found to be resistant to various viral and fungal infections (Wang et al., 1998; Lodge et al., 1993). The anti-viral activity of PAPs against human viruses has been described against Japanese encephalitis virus (Ishag et al., 2013), human immunodeficiency virus-1 (HIV-1) (Rajamohan et al., 1999), human T-cell leukemia virus-1 (HTLV-1) (Mansouri et al., 2009), herpes simplex virus (HSV) (Aron and Irvin, 1980), influenza (Tomlinson et al., 1974), hepatitis B virus (HBV) (He et al., 2008), and poliovirus (Ussery et al., 1977). PAPs low to moderate cytotoxicity to non-infected cells, in contrast to infected cells, makes PAPs very attractive candidates in the development of potential therapeutics and as protective agents against pathogens in transgenic plants.

Ricin is expressed in the seeds of the castor oil plant (R*icinus communis*) and is one of the most potent type II RIPs. It is highly toxic to mammalian cells as its A chain can efficiently be delivered into the cytosol of cells through the mechanism of its B chain. The B chain serves as a galactose/N-acetylgalactosamine binding domain (lectin) and is linked to the A chain via disulfide bonds (Lord et al., 2003). Ricin can induce 50% apoptosis in mammalian cells at concentrations below 1 ng/mL while showing no to low activity on plant and *E. coli* ribosomes. It is important to note however that the ricin A chain (RTA) on its own has less than 0.01% of the toxicity of the native protein in a cell culture test system. It was furthermore shown that RTA alone had no activity on non-infected and tobacco mosaic virus (TMV)-infected tobacco protoplasts alike. RTA lacks the ability to enter the cell without the action of the B chain (Watanabe et al., 1997). RTA depurinates a universally conserved adenine residue within the sarcin/ricin loop (SRL) of the 28S rRNA to inhibit protein synthesis. Though there are currently no commercially available therapeutic applications, RTA is extensively studied in the development of immunotoxins (Gilabert-Oriol et al., 2014).

The therapeutic potential of PAP and RTA has been explored for over thirty years, though side effects have limited clinical application. These proteins have shown very low cytotoxicity to non-infected cells; however, PAPs administration in mouse models has resulted in hepatic, renal and gastrointestinal tract damage with a median lethal dose (LD50) as low as 1.6mg/Kg (Benigni et al., 1995). Interestingly, RTA shows no toxicity even at high doses with similar half-life times. Nevertheless, all RIPs show immunosuppressive effects to various degrees. Many studies have described the various dose-limiting side effects of these proteins when used as immunotoxins (i.e. vascular leak syndrome, hemolytic uremic syndrome and pluritis, among others) (Schindler et al., 2011; Meany et al., 2015). Nonetheless, some patients achieved complete or partial remission against Refractory B-Lineage Acute Lymphoblastic Leukemia with sub-toxic dosages, for example.

Fusion and hybrid proteins of RTA and PAPs have also been developed in pursuit of selectively targeting infected cells and selectively recognizing viral components, though with limited success (Rothan et al., 2014; Chaddock et al., 1996). Indeed, the engineering of novel therapeutic fusion proteins with higher specificity, selectivity, and potency with fewer side effects is a leading strategy in drug development that is more often than not limited by our still new understanding of protein structure and function. Another limiting factor is the availability of efficient protein structure prediction and simulation software. It is only with the recent advent of more sophisticated *in silico* tools that protein engineering became a viable alternative to conventional drug discovery techniques.

The purpose of this research is to further characterize previously created and functional novel fusion proteins between RTA and PAPs (Hassan and Ogg, 2016, 2017). Here, we describe the development of an effective and scalable production system in *Escherichia coli* and of purification methods that enabled accurate determination of RTA-PAPs protein synthesis inhibition *in vitro*. We similarly describe the chimeric protein’s reduced cytotoxicity and significant anti-HBV activity by detecting HBV DNA in the supernatant, in HepAD38 cells, *in vitro*. The optimization of the protein design for maximum effect is also described using the most up-to-date protein structure and function prediction software available online and the resulting gain of function confirmed in a protein synthesis inhibition system *in vitro*.

## 2. Materials and Methods

### 2.1. E. coli in vivo expression system and Rabbit Reticulate Lysate protein synthesis inhibition

#### 2.1.1. Design of the DNA sequences of the proteins for *E. coli in vivo* expression system

The two cDNA coding for RTA-Pokeweed antiviral protein isoform 1 from seeds (RTA-PAPS1, 541 amino acids) and for RTA mutant-PAP isoform 1 from leaves (RTAM-PAP1, 556 amino acids including the N terminal 6-His tag) were optimized for *E*. coli expression and chemically synthesized by AscentGene.

#### 2.1.2. *E. coli in vivo* expression vector

The cDNA coding for RTA-PAPS1 and RTAM-PAP1 sequences described above were subcloned in *E. coli* pET30a expression vector (Novagene) to generate the pET30a-RP1 and pET30a-6H-RPAP1 vectors respectively. The inserts were validated by DNA sequencing.

#### 2.1.3. *E. coli in vivo* protein production

The above described vectors were transformed into *E. coli* BL21(DE3) cells (NEB) and expression of the proteins were examined from individual clones and analyzed by either Western blot using a monoclonal antibody specific to ricin A chain (ThermoFisher, RA999) or SDS gel stained with Comassie blue (ThermoFisher). Optimal conditions were determined and protein production induced in the presence of 1mM IPTG from 1L culture for each protein. The bacteria were then harvested by centrifugation, followed by lysing the cell pellets with 50ml of lysis buffer (50mM Tris-Cl, 150mM NaCl, 0.2% Triton X100 and 0.5mM EDTA). After sonication (3x2min), the soluble lysates were recovered by centrifugation at 35K rpm for 40min. The insoluble pellets were further extracted with 40ml of 6M Urea and the inclusion bodies (IB) were recovered by centrifugation at 16K rpm for 20min. Clarified IB were then dissolved with 20ml of buffer 8b (proprietary formulation of AscentGene). The soluble proteins were then recovered by centrifugation (please contact the authors for more details).

#### 2.1.4. *E. coli* protein purification

Ricin-PAPS1 proteins were purified by gel filtration column (Superdex 200 from GE Healthcare) under denaturing condition (6M Urea). Peak fractions were pooled and powder Guanidine was added to 5M for complete denaturing. Denatured Ricin-PAPS1 was then refolded by adding dropwise into the refolding buffer (50mM Tris-Cl, pH8.1, 0.4M L-Arginine, 0.5mM oxidized glutathione and 5mM reduced glutathione). After stirred at room temperature for 10min, the refolding reaction was further carried out at 4°C for >20hrs. Clarified and refolded Ricin-PAPS1 proteins were then concentrated before going through endotoxin removal and ammonium sulfate precipitation. The resulting mixture was dialyzed in the formulation buffer containing 20mM HEPES-Na, pH7.9, 20% glycerol, 100mM NaCl, 2.5mM TCEP and 1mM EDTA.

The purification of the native RTAM-PAP1 from soluble lysate was achieved by affinity versus Histag on Ni-sepharose column (GE Healthcare). After extensive washes with the lysis buffer, loosely bound proteins were eluted with the lysis buffer containing 40mM Imidazole (I40). RTAM-PAP1 proteins were eluted with elution buffer (20mM Tris-Cl, pH7.9, 100mM NaCl, 1mM EDTA and 300mM Imidazole). A second purification step using Hydroxylapatite column (GE Healthcare) was used to further separate RTAM-PAP1 from co-purified host proteins. A third purification step, gel filtration on a fast protein liquid chromatography (FPLC) column of Superose 12 (GE Healthcare), was necessary to completely get rid of degraded or/and premature protein products. The resulting mixture was dialyzed in the formulation buffer containing 20mM HEPES-Na, pH7.9, 200mM NaCl, 0.2mM CaCl2 and 0.5mM EDTA.

#### 2.1.5. Rabbit Reticulate Lysate protein synthesis inhibition

The inhibitory activities of RTA-PAPS1 and RTAM-PAP1 were tested by using the Rabbit Reticulate Lysate TnT^®^ Quick Coupled Transcription/Translation System and the Luciferase Assay System (Promega). Briefly, each transcription/translation reaction was performed according to the instructions for use (IFU) in the presence of a T7 Luciferase reporter DNA, and the Luciferase expression level was determined with a Wallac Microplate Reader. Transcription/translation runs were done twice with and without addition of five different concentrations of RTA-PAPS1 and RTAM-PAP1 in order to determine the inhibitory effect of the proteins. RTA-PAPS1 and RTAM-PAP1 concentrations were adjusted by taking sample purity into consideration.

### 2.2. Anti-HBV Assay

The anti-HBV assay was performed as previously described (Min et al., 2017) with modifications to use HepAD38 cells. ImQuest BioSciences developed a multi-marker screening assay utilizing the HepAD38 cells to detect proteins, RNA, and DNA intermediates characteristic of HBV replication. The HepAD38 cells are derived from HepG2 stably transfected with a single cDNA copy of hepatitis B virus pregenomic RNA, in which HBV replication is regulated by tetracycline. Briefly, HepAD38 cells were plated in 96-well flat bottom plates at 1.5 × 10^4^ cells/well in Dulbecco’s modified Eagle’s medium supplemented with 2% FBS, 380 μg/mL G418, 2.0 mM L-glutamine, 100 units/mL penicillin, 100 μg/mL streptomycin, and 0.1 mM nonessential amino acids (ThermoFisher). After 24h, six ten fold serial dilutions of RTA-PAPS1 prepared in the same medium were added in triplicate. Lamivudine (3TC from Sigma Aldrich) was used as the positive control, while media alone was added to cells as a negative control (virus control, VC). Three days later, the culture medium was replaced with fresh medium containing the appropriately diluted RTA-PAPS1. Six days following the initial administration of RTA-PAPS1, the cell culture supernatant was collected, diluted in qPCR dilution buffer, and then used in a real-time quantitative qPCR assay using a Bio-Rad CFX384 Touch Real-Time PCR Detection System. The HBV DNA copy number in each sample was interpolated from the standard curve by the supporting software. A tetrazolium dye uptake assay (ThermoFisher) was then employed to measure cell viability, which was used to calculate cytotoxic concentration (TC50).

### 2.3. Protein Design Optimization

#### 2.3.1. Physiochemical profiling and specific structural features

The molecular profile of the protein was determined using Protparam tool of ExPASy (Gasteiger et al., 2005), and the solubility of these proteins was determined using Predict Protein (Yachdav et al. 2014). The presence of disulfide bonds was determined using DiANNA 1.1 webserver (Ferre and Clote, 2005a, 2005b, 2006). Functional effects of point mutations were determined using SNAP2 of Predict Protein.

#### 2.3.2. Structure modeling

The structure of the protein was predicted by fold recognition methodology using I-TASSER (Zhang, 2009; Roy et al., 2012; Yang and Zhang 2015) and Phyre2 (Kelley et al., 2015) prediction server. The determined protein structures were then validated by Verify 3D (Bowie et al., 1991; Lüthy et al., 1992). The quality of the structure was determined using QMEAN6 program of the SWISS-MODEL (Benkert et al., 2008) workspace.

#### 2.3.3. Design of RTAM-PAP1

Three major changes were made to RTA-PAPS1 in order to increase its solubility, its efficacy against infected cells and to further reduce its toxicity.

Firstly, two points mutations of least effect on function as predicted by SNAP2 of Predict Protein were introduced into the RTA moiety to replace the Cysteine (Cys) residues with Alanine residues in order to completely avoid unwanted disulfide bonds formation at position 171 and 259 (C171A and C259A) to create RTA mutant RTAM.

Secondly, the natural semi-flexible linker previously used was replaced with a newly designed soluble flexible G rich linker with a rigid CASP2 recognition site (GGGGSDVADI(GGGGS)2) to allow better autonomous function of each moiety with minimal steric hindrance and to further enhance the chimeric protein’s ability to induce cell apoptosis (Chen et al., 2013).

Thirdly, A different variant than PAPS1 was used, PAP1, retrieved from National Centre for Biotechnology Information database (NCBI) with access number **<u>P10297.2</u>** in order to further enhance activity against HBV and further reduce toxicity of the chimeric protein.

Lastly, a 6-His tag was added at the N terminal of the protein RTAM-PAP1 in order to minimize effect on structure and function and to increase native protein recovery from *E. coli* production.

## 3. Results

### 3.1. Production and Purification of recombinant RTA-PAPS1 in E. coli culture

The production of fusion Ricin A Chain-Pokeweed Antiviral Protein from Seeds Isoform 1 (RTA-PAPS1) in *E. coli* was found to be significantly better at 30°C than at 37°C (results not shown). In order to optimize the amount of protein produced from 1L at 30°C, three media were tested: M9 (M9), Luria Bertani (LB) and terrific broth (TB). Soluble lysate (Sol) and inclusion body (IB) from each sample were analyzed by SDS PAGE and visualized by Coomassie blue staining (Figure 1.a). As can be seen, almost all of the overexpressed RTA-PAPS1 proteins were in the form of inclusion bodies, which were almost completely insoluble in either 6M Urea or 6M Guanidine. A total of 28 buffers were tested and only the denaturing buffer 8b (proprietary formulation of AscentGene) was able to dissolve more than 50% of the Ricin-PAPS1 present in the inclusion bodies. Once the soluble proteins were recovered and purified through gel filtration column Superdex200 (single step) in their denatured form, they were allowed to refold for over 20hrs in a refolding buffer before being concentrated. The resulting protein was found to be at a concentration of 0.22mg/ml at >90% purity (Figure 1.b), which was the best result we obtained thus far after testing numerous production methods and bacterial strains, as previously described (Hassan and Ogg, 2016, 2017).

**Figure 1.**
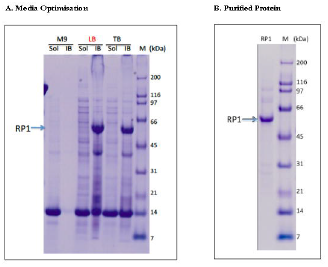
Medium Optimization and Protein Purification. a) Medium optimization for Ricin-PAPS1 (RP1) expression. Three different growth media including M9 (M9), Luria Bertani (LB) and terrific broth (TB) were tested for Ricin-PAPS1 expression at30°C. Soluble lysate (Sol) and inclusion body (IB) from each sample were analyzed by SDS PAGE and visualized by Coomassie blue staining, b) Validation of purified Ricin-PAPS1 protein. Recombinant Ricin-PAPS1 was produced in 1L of culture that was induced with the optimized condition (LB medium with 1mM IPTG at 30°C for 4hrs) and purified from inclusion bodies through gel filtration before refolding, concentration and dialysis. The resulting protein of approx. 60.5kDa was >90% purity determined by SDS-PAGE.

### 3.2. Inhibitory activity of recombinant RTA-PAPS1 in the Rabbit Reticulate Lysate TnT^®^ system

The inhibitory activity of RTA-PAPS1 was determined using 5 different concentrations of purified RTA-PAPS1 in duplicate on the Rabbit Reticulate Lysate TnT^®^ system using Luciferase as control. A Luciferase assay was used to determine Luciferase expression levels using a luminometer. The resulting plot is shown in Figure 2. As can be observed, RTA-PAPS1 has an IC50 at 0.06nM, slower than RTA IC50 at 0.03nM but comparable to PAPS IC50 at 0.07 (Hassan and Ogg, 2017; Poyet and Hoeveler, 1997; Hale, 2001). The IC100 however is attained faster than all of them at 0.24nM for RTA-PAPS1 while RTA IC100 is 0.60nM. These results show that RTA-PAPS1 is bioactive with a synergetic activity between the RTA and PAPS1 moieties being noticeable.

**Figure 2.**
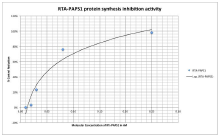
Test of purified RTA-PAPS1 in the TnT transcription/translation assay. Five different concentration points (0.01 nM, 0.02nM, 0.03nM, 0.08nM, 0.25nM) were examined. Values are calculated as percent inhibition of Luciferase protein synthesis compared to control. Results represent the mean for two individual experiments and the curve is the logarithmic regression.

### 3.3. Anti-HBV evaluation of recombinant RTA-PAPS1 in HepAD38 cells

Recombinant RTA-PAPS1 was evaluated for anti-HBV activity and cytotoxicity in the HBV chronically infected cell line AD38 using a six concentrations dose response assay in triplicate. The lamivudine (3TC) control compound was evaluated in parallel. The antiviral efficacy based on quantified DNA copies in the supernatant of both compounds are shown in Figure 3 in a plot form. RTA-PAPS1 yielded a half-maximal response concentration value (EC50) of 0.03nM while 3TC yielded an EC50 of 0.3nM, which is a ten-fold difference. RTA-PAPS1 was not cytotoxic to HepAD38 cells at concentrations up to 600nM. These results led to a therapeutic index for RTA-PAPS1 of >21818, which is a huge improvement over values given in literature (EC50 of 330nM and a therapeutic index of 9.3 for PAPS1 alone under comparable conditions on HepG2 2.2.15 cells) (He et al., 2008). These results clearly show the significant anti-HBV activity of RTA-PAPS1.

**Figure 3.**
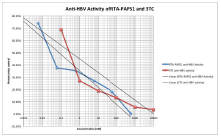
Anti-HBV evaluation of RTA-PAPS1. Recombinant RTA-PAPS1 was tested for its anti-HBV activity using 6 concentrations using a serial dilution by a factor of 10 in growth media (600nM, 60nM, 6nM, 0.6nM, 0.06nM, 0.006nM for RTA-PAPS1 and 10000nM, 1000nM, 100nM, 10nM, 1nM, 0.1nM for 3TC). Values are calculated as percent of virus DNA control [(amount of virus DNA in treated sample/amount of virus DNA in untreated sample) x100]. Results represent the mean for three individual experiments.

### 3.4 Protein Design Optimization

The design of the recombinant protein RTA-PAPS1 was completely revisited in order to further enhance the effect of the chimeric protein on HBV, reduce general toxicity and increase solubility to improve expression. The resulting design Ricin A Chain Mutant-Pokeweed Antiviral Protein from Leaves (RTAM-PAP1) was run through I-Tasser and Phyre2 and the resulting 3D models validated by Verify 3D. The model generated by Phyre2 passed Verify 3D while the one generated by I-Tasser failed. The one generated by Phyre2 was thus chosen as one of the templates to run I-Tasser again. The newly generated structure by I-Tasser scored higher on Verify 3D than the one generated by Phyre2 and was thus chosen as the model for the other software. The proper disulfide bond formations were confirmed by DiANNA 1.1 webserver (at positions 328–553 and 379–400). The new model had a normalized QMEAN4 score of >0.6 and the introduction of the rigid CASP2 recognition site into the flexible linker at position 280-285 insured safe distance between the two proteins to safeguard the function of both moieties and minimize steric hindrance as can be seen in Figure 4. The grand average of hydropathicity was reduced from -0.236 for RTA-PAPS1 to -0.265 for RTAM-PAP1 as was determined by ProtParam, which represents an improvement of 12% in hydrophilicity (results not shown).

**Figure 4.**
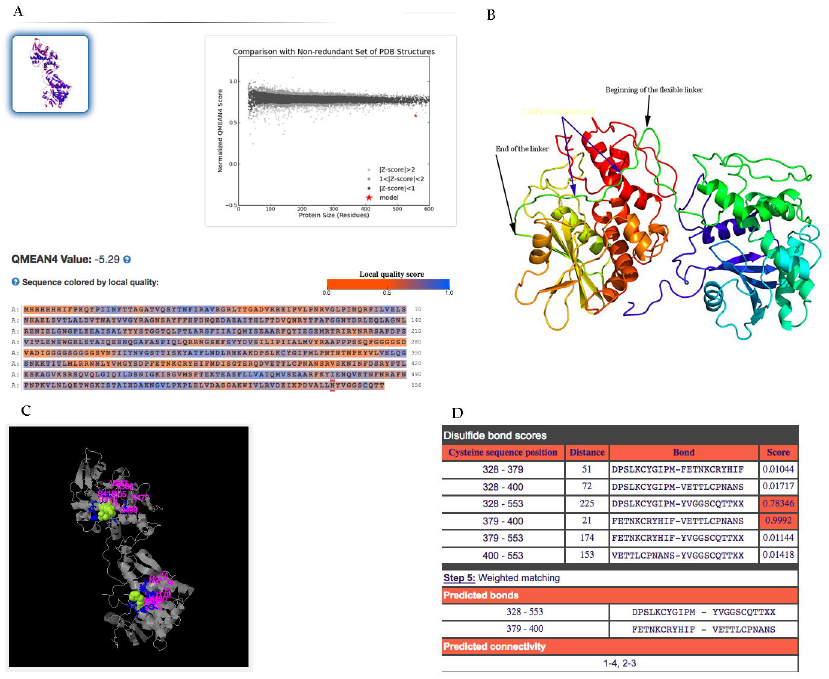
Predicted 3D Protein Structure. a) QMEAN4 results showing a normalized value >0.6 with the protein sequence in the bottom and the model in the up left corner. b) Protein structure as determined by Phyre2 with the black arrows showing the flexible linker at position 275–294 and the blue arrows showing the CASP2 recognition site at position 280–284. Image colored by rainbow (blue to red) N → C terminus. c) The ligand binding sites of RTAM moiety (up) and of PAP1 moiety (down) as determined by I-Tasser (using the Phyre2 model as one of the templates), d) The predicted disulfide bonds at 328–553 and 379–400 by DiANNA 1.1 webserver.

### 3.5. Production and Purification of recombinant RTAM-PAP1 in E. coli culture

The production of RTAM-PAP1 was first tested under the same conditions as previously determined for RTA-PAPS1 and resulted in good production of native proteins. Soluble RTAM-PAP1 was recovered from the lysate, purified by Ni-sepharose column and analyzed by SDS-PAGE and Western Blot (Figure 5). The production from 1L culture under the same conditions gave equally good results (Figure 6a.). The purified proteins were then submitted to a second purification step using hydroxylapatite column, which showed good separation of RTAM-PAP1 from co-purified host proteins (Figure 6b). The degraded (or premature) products were further separated by gel filtration on an FPLC column of Superose 12 (Figure 7a) and the purest fraction (F15) reached >95% homogeneity at a concentration of 0.1mg/ml (Figure 7b) and was used for the protein synthesis inhibition assay.

**Figure 5.**
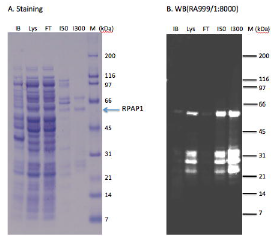
Test Production and Purification of native RTAM-PAP1. a) Loosely bound proteins were washed with the lysis buffer containing 50mM imidazole (150) on a Nisepharose column and RTAM-PAP1 (RPAP1) proteins were then eluted with the elution buffer containing 300mM Imidazole (I300). b) The Western Blot using ricin a chain antibody RA999 confirmed the presence of RTAM-PAPS1 at approx. 61.5kDa. The bands between 21kDa and 32kDa are assumed to be degraded or/and premature RTAM-PAP1 proteins.

**Figure 6.**
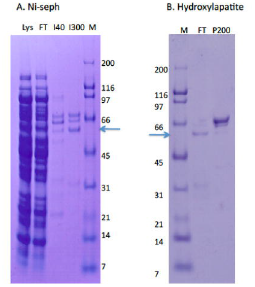
2 steps Purification of RTAM-PAP1 from soluble lysate (Lys) from 1L culture. a) Loosely bound proteins were washed with the lysis buffer containing 40mM imidazole (I40) on a Ni-sepharose column and RTAM-PAP1 proteins were then eluted with the elution buffer containing 300mM Imidazole (I300). b) Co-purified host cell proteins were further separated by a hydroxylapatite column. Most RTAM-PAP1 proteins were retained in the flow through (FT) fraction, while most host cell proteins were bound to the hydroxylapatite column (P200 elution).

**Figure 7.**
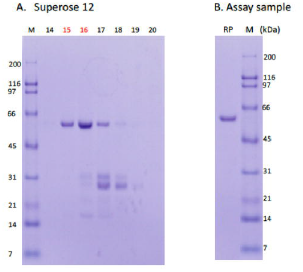
Third purification step of RTAM-PAP1 by Superose 12 column. a) RTAM-PAP1 was peaked at fraction 15 and 16. The purest fraction (F15) was estimated at >95% homogeneity b) and was used for the inhibition assay.

### 3.6. Inhibitory activity of recombinant RTAM-PAP1 vs. RTA-PAPS1 in the Rabbit Reticulate Lysate TnT^®^ system

The inhibitory activity of RTAM-PAP1 was determined using 5 different concentrations in duplicate of purified RTAM-PAP1 on the Rabbit Reticulate Lysate TnT^®^ system using Luciferase as the control as previously described. The resulting comparative plot of the activity on protein synthesis of both fusion proteins is shown in Figure 8. As can be observed, RTAM-PAP1 has an IC50 at 0.03nM, the same as RTA IC50 at 0.03nM and twice as fast as RTA-PAPS1 IC50 at 0.06nM and about ten times faster than PAP1 IC50 at 0.29nM (Poyet et al., 1997). The IC100 however is attained faster than all of them at 0.07nM for RTAM-PAP1 while RTA IC100 is 0.60nM, PAP1 IC100 at 1.1nM and RTA-PAPS1 IC100 at 0.24nM. These results show that RTAM-PAP1 is bioactive, both moieties’ complementary catalytic activities functional, with minimal steric hindrance if any, and with a significant gain of function.

**Figure 8.**
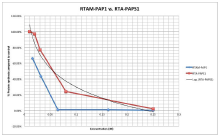
Comparative inhibition activity of RTAM-PAP1 and RTA-PAPS1 in the TnT transcription/translation assay. Five different concentration points (0.01 nM, 0.02nM, 0.03nM, 0.08nM, 0.25nM for RTA-PAPS1 and 0.02nM, 0.03nM, 0.06nM, 0.16nM, 0.40nM for RTAM-PAP1) were examined. Values are calculated as percent Luciferase protein synthesis compared to control. Results represent the mean for two individual experiments and the curve is the logarithmic regression for RTA-PAPS1.

## 4. Discussion

The chimeric protein RTA-PAPS1 was expressed only in inclusion bodies with very little solubility, except under heavy denaturing conditions. The refolding process was successful to a certain extent as more than one conformation was observed. This was probably due to the two free Cysteine residues in RTA and the close proximity of Cys at position 260 to the Cys residues at position 364 and 385 (confirmed by DiANNA 1.1 webserver and I-Tasser). The addition of TCEP was necessary and a difference in bioactivity (>2 fold) was observed between conformations (results not shown). RTA-PAPS1 was nonetheless very bioactive and with a certain degree of synergetic activity between RTA and PAPS1, probably limited by steric hindrance due to the semi-rigid quality of the linker. This was confirmed during the anti-HBV assays. The significant anti-HBV activity was apparent and probably due to the ability of both moieties to depurinate rRNA but also polynucleotide, single-stranded DNA, double stranded DNA and mRNA (He et al. 2008; Ling et al. 1994). HBV is a double stranded DNA reverse transcriptase virus. A high standard deviation was observed between samples at the same concentration for anti-HBV activity and dubious results were seen at the highest concentration of 600nM for cytotoxicity. This is probably due to the presence of different conformations of the same protein and a high protein buffer solution ratio to growth media at 600nM (>30%) respectively. Furthermore, it was observed that the steric hindrance diminished the ability of PAPS1 to penetrate infected cells by >2 fold as the activity of RTA-PAPS1 on other viruses such as HIV-1 for example was much lower than that of PAPS1 alone (results not shown). For these reasons, the decision to redesign the fusion protein with PAP1 and a new linker was taken.

The fusion protein RTAM-PAP1 expression went very well as we almost exclusively obtained native protein production with high solubility (barely any in inclusions bodies). It was however necessary to use a three step purification protocol in order to obtain soluble proteins with >90% homogeneity. Nonetheless, 0.1mg of protein at >95% purity and 0.22mg of protein at >90% purity were obtained from 1L of culture. This yield is probably explained by the increased toxicity of PAP1 to *E. coli* compared to that of PAPS1 (>10 fold) (Honjo et al., 2002). The bioactivity of RTAM-PAP1 was increased, much more than expected with very little to no sign of steric hindrance. The introduction of the two point mutations in the RTA moiety and of the flexible linker really made a difference in solubility and activity. Also, perhaps, finetuning the formulation buffer to better preserve protein integrity allowed for optimum activity. The synergetic effect of both moieties was very apparent and probably due to the fact that RTA and PAP1 do not dock onto the ribosome at the same site and, thus, led to a reduction of partially depurinated and still functional ribosomes (Chaddock et al. 1996).

## 5. Conclusion

The chimeric protein between RTA and PAPs are potent novel anti-viral proteins with gain of function in protein synthesis inhibition activity and anti-HBV activity *in vitro* with minimal cytotoxicity. The introduction of two point mutations in RTA and of a flexible linker greatly improved solubility and activity. RTAM-PAP1 can be overexpressed, recovered and purified from soluble lysate. It is expected that the anti-viral properties of RTAM-PAP1 against plant and animal pathogens will be even greater than that of either RTA-PAPS1 or PAPs with even lesser general toxicity. It is the opinion of the authors that a full characterization of RTAM-PAP1 activity against chronic HBV infection be done both *in vitro* and *in vivo* as it is a potential potent new therapeutics to be used as a standalone or in combination with existent therapies.

**Availability of Data and Materials:** All data generated or analyzed during this study are included in this published article, further details or raw data are available from the corresponding author on reasonable request.

**Conflicts of Interest:** “This work was funded by Ophiuchus Medicine which owns the rights to the patent pending of the fusion proteins described in this study. The authors are either directly or indirectly affiliated to Ophiuchus Medicine.”

## Footnotes

i **Abbreviations:** Ricin A Chain (RTA), Pokeweed antiviral proteins (PAPs), Pokeweed antiviral protein isoform 1 from seeds (PAPS1), Pokeweed antiviral protein isoform 1 from leaves (PAP1), 50% protein synthesis inhibitory concentration (IC50), 100% protein synthesis inhibitory concentration (IC100) and half-maximal response concentration of a drug (EC50).

